# Nutrition and density dependence of spontaneous female-biased dispersal in *Drosophila melanogaster*

**DOI:** 10.1101/2024.05.27.596025

**Authors:** Subhasish Halder, Utkarsh Bhore, Bodhisatta Nandy

## Abstract

Dispersal is often essential for the attainment of Darwinian fitness, especially for species living on spatially structured, heterogeneous habitats. Theoretically, sex-specific resource requirement can drive the two sexes to disperse differently, resulting in sex biased dispersal (SBD). Understanding ecological factors affecting SBD is important. Using an experimental two-patch dispersal setup we measured spontaneous dispersal in laboratory adapted populations of *Drosophila melanogaster* under a set of common, interlinked ecological scenarios relating to – (a) dietary ecology and (b) adult density. We found deteriorating overall nutritional quality of food affects strength of SBD, and female dispersal is particularly sensitive to availability of protein. Adult density had sex specific effect on dispersal. Female dispersal was found to be density independent but males showed increased dispersal at higher density. Female tend to disperse more from male biased patch likely to avoid male harassment whereas absence of female drives male dispersal solidifying mate-finding dispersal hypothesis. These evidences of dispersal suggest that variation in dietary ecology and intraspecific competition can affect the degree and strength of existing SBD and thereby male-female interactions in a patch potentially affecting fitness components and population dynamics.

## Introduction

Most naturally occurring populations show spatial structure such that a population can be viewed as a set of sub-populations distributed across various habitable patches or sub-populations. The ability to disperse across these patches – accessing the rich patches, and avoiding the poor ones, can essentially be a determinant of fitness (Bowler & Benton, 2005; Travis et al., 2013). Importantly, dispersal is key for most species to survive in the dynamic natural environment as it allows exploitation of spatially and temporally variable resources (Ronce, 2007; Clobert, 2009; Reyes, 2023). From protists to vertebrates, resource limitation in a patch increases dispersal (Fronhofer et al., 2017). Within a species, spatiotemporally variable resources can favour evolution of dispersal genotypes aiding in adaptation to a novel ecological setup (Friedenberg, 2003). Causes and constraints in the evolution of dispersal traits is an important yet poorly understood area in Evolutionary Ecology. Interestingly, many sexually reproducing animals show sex difference in their tendency to disperse. Ecological determinants of sex biased (either male-biased or females biased) dispersal, and its eco-evolutionary consequences is central to adaptation under spatially structured habitat.

In polygamous species, owing to differences in reproductive investment, including parental investment, resources that limit fitness are often different in the two sexes (Li & Kokko, 2019). Female fitness largely depends on feeding, nesting and/or breeding ground. Whereas for males, especially in species with conventional sex roles, fitness is determined by the number of mates and paternity success. Such sex-specific fitness interest can lead to the evolution of sex biased dispersal (SBD). Literature on SBD supports ‘resource competition hypothesis’ which indicates that there are two ecological factors that are responsible for sex difference in dispersal - local resource defense and local mate competition (Greenwood, 1980). The latter being supported by the observation that polygyny probably favours male-biased dispersal (for example, in mammals, Mabry et al., 2013; Hooft et al., 2018 ), while monogamy favours female-biased dispersal (for example, in birds, Clarke et al., 1997; Wolff et al., 1998). Beyond mammals and birds, SBD have been increasingly reported in other taxa, and in diverse mating systems (Trochet et al., 2016). Interestingly within a taxon there are examples of both male-biased and female-biased dispersal (Li & Kokko, 2019). There is at least some evidence suggesting SBD to be condition dependent (Mishra et al., 2018; Bitume et al., 2013). Further, being energetically expensive, behaviour and physiology of dispersal is also likely to be constrained by life-history constraints, including overall availability of metabolic resources (Stevens et al., 2012; Venkateswaran et al., 2017). Therefore, within the paradigm of local resource competition, it is important to understand the ecological factors generating diversity in dispersal tendency, both within and across species.

Dispersal is often associated with a suit of correlated traits, collectively known as dispersal syndrome, which includes resource extensive behaviour and physiology such as, locomotion, exploration, aggression, higher metabolic rate etc. (Tung et al., 2018). Investment in these traits is expected to be competitive against life-history traits including longevity, and reproductive rate (Bonte et al., 2012). Sex difference in life history can generate sex biased dispersal (Clutton-Brock et al., 2002; Li & Kokko, 2019). For example, species where males are known to have live-fast-die-young life-history, males are often more dispersive (Jungwirth et al., 2023). Theoretically, the converse, i.e., sex difference in dispersal resulting from sex difference in life-history, is possible. However, this is unclear if sex difference in life-history itself can stem from sex difference in dispersal syndrome. As it turns out, the interconnection between dispersal syndrome and life-history is a relatively new research theme, with key results reported relatively recently (Tung et al., 2018).

Theories predict that the sex that pays more fitness cost from local competition should disperse more often (Perrin & Mazalov, 2000). Trochet et al., (2013), using an experimental setup that mimic interconnected-patch habitat, tested the theory of SBD by manipulating sex ratio in populations of the butterfly *Pieris brassicae*, but did not find SBD. A semi aquatic, flight capable insect *Notonecta undulata* shows male biased dispersal rate, however varying sex ratio did not affect the existing pattern (Baines et al., 2017). On the other hand, Mishra et al., (2018) showed density dependency of enforced dispersal (dispersal upon experimental starvation), and the resultant variation in the degree of SBD in fruit fly *Drosophila melanogaster* laboratory populations. Interestingly, female-biased dispersal was observed at a low adult density, and male-biased dispersal at a high-density condition (Mishra et al., 2018). However, it is not clear how the degree of spontaneous sex biased dispersal tendency change in the face of variation in the prevailing conditions including dietary ecology, density, and sex ratio, despite the fact that such variation is commonplace in any natural meta-population.

Here, we investigated this problem using laboratory populations of *D. melanogaster,* which has been a useful experimental model even for dispersal biology research (Dobzhansky and Wright 1943; Coyne et al., 1982, 1987; Tung et al., 2018a, 2018b; Dukas, 2020; Mishra et al., 2018, 2022). Importantly, most dispersal studies in *D. melanogaster* have used extreme starvation and/or desiccation driven movement of individuals to assess dispersal. Physiological condition can alone force individuals to disperse, independent of sex and other cues related to resources (Wang et al. 2014; Mishra et al. 2022). We refer to this mode of dispersal as ‘forced dispersal’ to differentiate this behaviour from the inherent tendency of individuals to move between habitat patches, which will be referred to as ‘spontaneous dispersal’. Previously, we have reported an experimental setup to study spontaneous dispersal using a two-patch experimental arena without the use of prior starvation/desiccation conditioning and have also shown female-biased dispersal pattern (Halder et al., 2024). Here, using this experimental setup, we investigated the role of two major ecological factors with putative role in modulating spontaneous dispersal – (a) diet (overall nutritional deprivation, carbohydrate and protein deprivation), (b) density (adult density and sex ratio). We hypothesised that female dispersal to be more sensitive to nutritional manipulation compared to males, while the latter sex to be more responsive to the intensity of mate competition.

## Material and methods

We used a set of outbred, laboratory-adapted populations of *Drosophila melanogaster* (BL_1-5_), maintained under a 14-day discrete generation cycle, 24 h light, at 25 °C (±1) temperature on standard banana-jaggery-yeast food medium (see supplementary information for food composition). Flies are grown in culture vials (25 mm diameter × 96 mm height) at a density of ∼70 per 6-8 ml food in each vial, forty such vials make up one population. On day12 post-egg collection, the adult flies are transferred into a population cage (dimension: 23×20×15cm) and thereafter maintained in the cage as a population of ∼2,800 individuals. On day14, oviposition substrate is provided and eggs are collected from window of 18 hours.

These eggs are cultured in fresh food vials following the above-mentioned density to start the next generation. The details of the history and maintenance of the populations can also be found in Dasgupta et al., 2022 and Halder et al., 2024. The five replicate BL populations have adapted to the above-mentioned maintenance regime for several hundred generations. The 18h window during which eggs are collected to start the next generation, corresponds to the populations’ Darwinian fitness window. Therefore, progeny produced during this window is an accurate measure of fitness, particularly in this population system. The following experiments were conducted with three replicate populations following the philosophy of statistical block design.

### Dispersal assays

Eggs were collected from a randomly chosen population (the assay was done with BL_3,_ BL_4_, BL_5_), and cultured in standard vials at a density of 70 per vial with ∼8ml food. The adults were allowed to emerge in these vials, upon which they were transferred to fresh food vials with standard food. On day12 from the egg-collection, i.e., at 2-3 days post-eclosion, two-patch dispersal assays were set up to measure dispersal tendencies of the experimental flies. This particular adult age i.e., 2-3 days post-eclosion was opted to assess spontaneous dispersal dynamics because it is the most relevant age for adult interaction and fitness in this model population system as discussed in the previous paragraph.

The dispersal assay setups used here were two-patch systems constructed by connecting a culture vial (with food) to a standard Drosophila culture bottle (Laxbro) with a tygon tube (Tarsons) of 0.6 cm inner diameter and 31cm length (see Figure 1). The vial was used as the source patch, where flies were introduced to start the assay, while the bottle is used as the sink. To ensure unidirectional movement of the flies, the hole in the bottle was fitted with a pipette tip, cut in such a way that it allowed flies to exit the tube through it and enter the bottle (see Figure 1). Observations in our laboratory showed that without the pipette tip arrangement, flies moved in both directions unabated (i.e., vial to bottle, as well as bottle to vial). However, this setup allowed only source to sink movement, but stopped all movement in the reverse direction.

**Figure 1:**
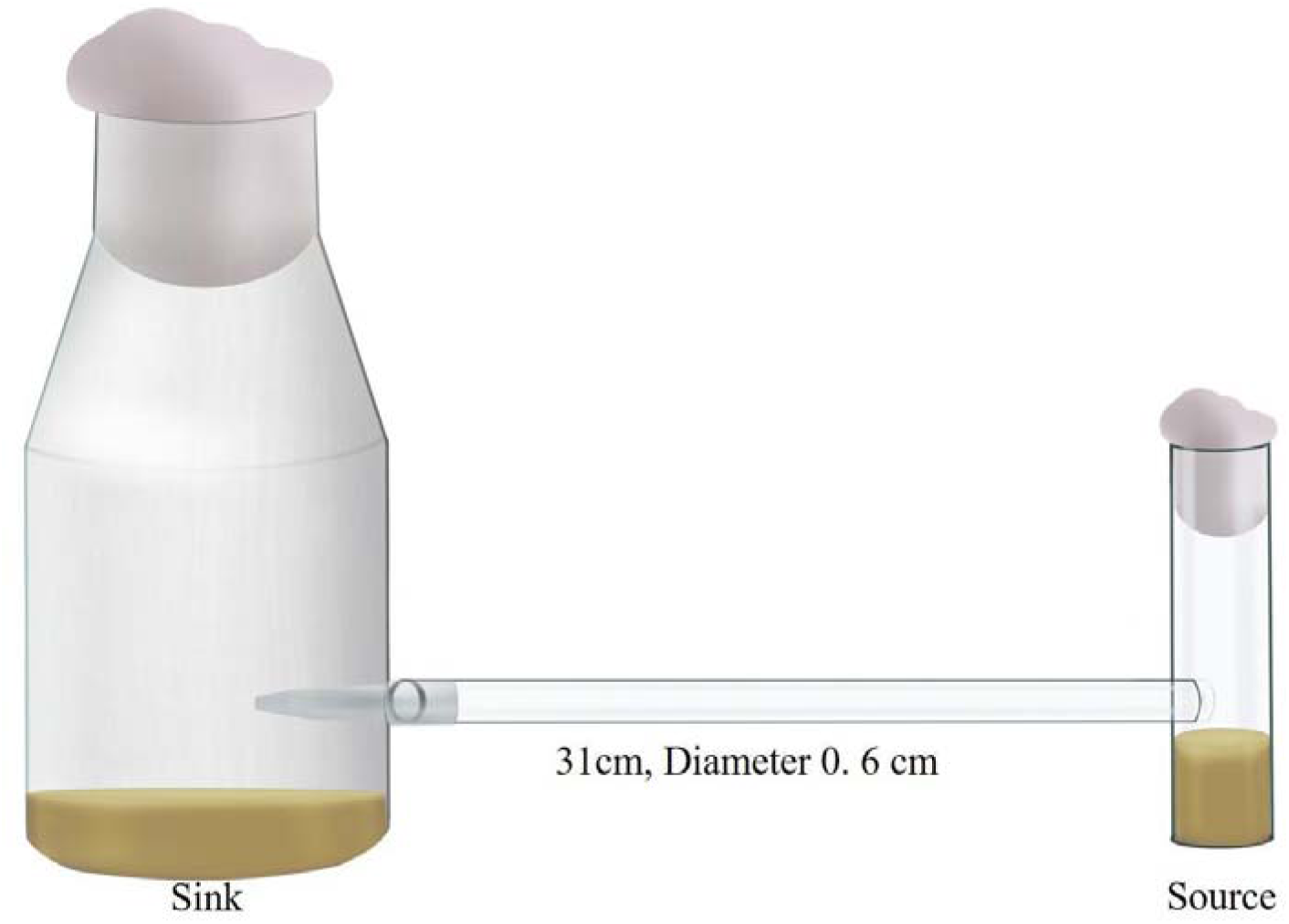
A two-patch setup for dispersal assay. A culture food vial (Source patch) connected to a standard Drosophila culture bottle (Sink patch) by a tube of 0.6 cm inner diameter and 31cm length.

To start the assay, number of males and female flies were counted by using light CO_2_- anaesthesia and introduced in a source patch (i.e., the vial). The setup was then placed horizontally in a well-lit place with illumination on all four sides. These setups were randomly placed to avoid any positional bias. They were left undisturbed for six hours for observation. Number of flies reaching the sink patch (i.e., the bottles) at the end of six hours were counted. Dispersal was quantified as the proportion-moved for each sex at the end of 6 hours. Based on our preliminary observations, by 6 hours, on an average more than 50% females move away from the source patch whereas males hardly move. To avoid the confounding effect of changing sex ratio, we considered proportion dispersed at 6 hours as our unit of analysis.

### Experiment 1: Effect of diet and nutrition on dispersal tendency

Reproduction and survival is the primary fitness component and often trades off under challenges related to diet and are therefore central to the life history of most organisms (Dasgupta et al., 2022). Heterogeneity in dietary ecology and nutrition shapes life history traits and hence poor diet can have detrimental fitness consequences (Chapman & Partridge 1996). Additionally, due to sex specific differences in resource allocation and nutrition requirements, two sexes can respond differently when faced challenges related to dietary ecology (Zajitschek et al., 2016; Li & Kokko 2018; Zajitschek et al., 2019). Therefore, by theory, available nutrition of a food patch can influence existing sex-biased dispersal dynamics, since female reproductive traits are also directly affected by dietary ecology. To address this issue, we conducted two assays.

Therefore, in the first assay (Experiment 1.1), we manipulated diet by diluting all the nutritional components of standard fly food and created a gradation of source patch diet treatments that varied in quality. The source patch treatment conditions were - Control or Standard food (100%), 75%, 50%, 25% of standard food and 0% that is only non-nutritive agar media (See Table S1 for the recipe). Following that, 30 adult flies with 1:1 sex ratio were introduced in the source patch and dispersal tendency was measured only for both the sexes.

Emerging evidence using geometric framework of nutrition suggests that specific macronutrients (and not mere diet or calorific value) trigger life history trade off, including maintenance of immunity related traits (Cotter et al., 2013; Rapkin et al., 2018). In addition, females of *Drosophila melanogaster* make important feeding and foraging decisions to optimize macronutrient uptake (Lihoreau et al., 2016; Camus et al., 2018). A logical extension of that could be - variation in the macronutrient such as protein and carbohydrate in diet can affect SBD pattern, especially the strength of female-biased dispersal. To test this prediction, in the second assay (experiment 1.2), we generated source food conditions that varied only in two main macronutrient - either protein (manipulated with yeast amount in food) or carbohydrate content (manipulated the amount of sugarcane jaggery, one of the sources of carbohydrate) generating treatment as follows-Control or Standard food, 25% protein of Control, Minimum-protein food, 25% carbohydrate of Control, Minimum - carbohydrate food (See Table S2 for the recipe). Following that, 30 adult flies in 1:1 sex ratio were introduced in the source patch and dispersal tendency was measured only for females.

### Experiment 2: Effect of adult density and sex ratio on dispersal tendency

Sex difference in dispersal can have an apparent impact on the number of available mates and competition in a patch and thus varying inter- and intra-sexual competitive scenarios across habitable patches. To cope with existing inter- and intra-sexual competition, organisms can not only adopt phenotypic plasticity but also resort to switching their behavioural strategies (Berglund, 1995; Jirotkul, 1999). Varying competition which inevitably results from the changes in demographic factor such as density and operational sex ratio can also affect reproductive output (Kokko & Rankin 2006) and facilitate sex-biased dispersal (Kokko & Rankin 2006; Li & Kokko 2018; Li & Kokko, 2019). Surprisingly, there are insufficient studies using controlled experimentation which specifically investigate SBD under mentioned context (however see, Trochet et al., 2013; Mishra et al., 2018). We conducted two separate assays to investigate (a) density effect (under equal sex ratio) of spontaneous dispersal, and (b) the effect of sex ratio on spontaneous dispersal tendency in the two sexes.

In the first assay (experiment 2.1), we introduced adult flies in the source patch with three different holding densities – Standard or control (70 flies of equal sex ratio), Low density (34 flies of equal sex ratio) and High density (210 flies of equal sex ratio) and quantified spontaneous dispersal tendency. Sex ratio and food quality were kept standard for the assay.

In the second assay (experiment 2.2), we tested the effect of adult sex ratio on dispersal tendency. To this effect, 32 adult flies were introduced in source patch of standard food with varying operational sex ratio – Standard or equal sex ratio (referred as ES), Male biased (24 males and 8 females or 3:1 ratio, referred as MB), Female biased (8 males and 24 females or 1:3 ratio, referred as FB), All male group (32 males, referred as AM), All female group (32 females, referred as AF). Adult density and food quality were also kept standard for the assay.

### Statistical analysis

All experiments were conducted using three randomly picked populations (i.e., BL_3_, BL_4_ and BL_5_ populations) as independent blocks and with 12 replicates per source patch treatment condition per block. Proportion dispersed was analysed using General Linear Mixed Model (GLMM) - using package `lme4` and function `glmer` with binomial family distribution and logit link function in R (R Version 4.2.0). Throughout, block and interactions involving block were treated as random factors (see Table S7 for the detailed model). Treatment (diet and Sex ratio) were considered as fixed factors. In the experiment 2.2, proportions dispersed were analyzed for two sexes separately taking sex ratio treatments as fixed effect. Following the analysis of variance using `Anova` function (package `car).` Post-hoc pairwise comparisons were done with `pairwise` function in package `emmeans`.

## Results

### Experiment 1

Analysis of dispersal data from the first assay (Experiment 1.1) revealed significant effects of Treatment, Sex and Treatment × Sex (Table 1, Figure 2). For females, proportion dispersed was progressively higher with increasing dilution. However, multiple comparisons indicated that the extreme dilution (i.e., 0%) showed significantly higher proportion dispersed compared to the rest of the levels. Two intermediate levels – i.e., 75% and 50%, showed intermediate values, but these were not significantly different from either 0% or the lower dilutions (i.e., control and 25%). For males, while on an average there was much less dispersal compared that observed in females, the proportion dispersed showed little change from control up to 75%. Only at 0%, i.e., under virtual starvation, there was a 2-3 fold jump in proportion dispersed.

**Figure 2:**
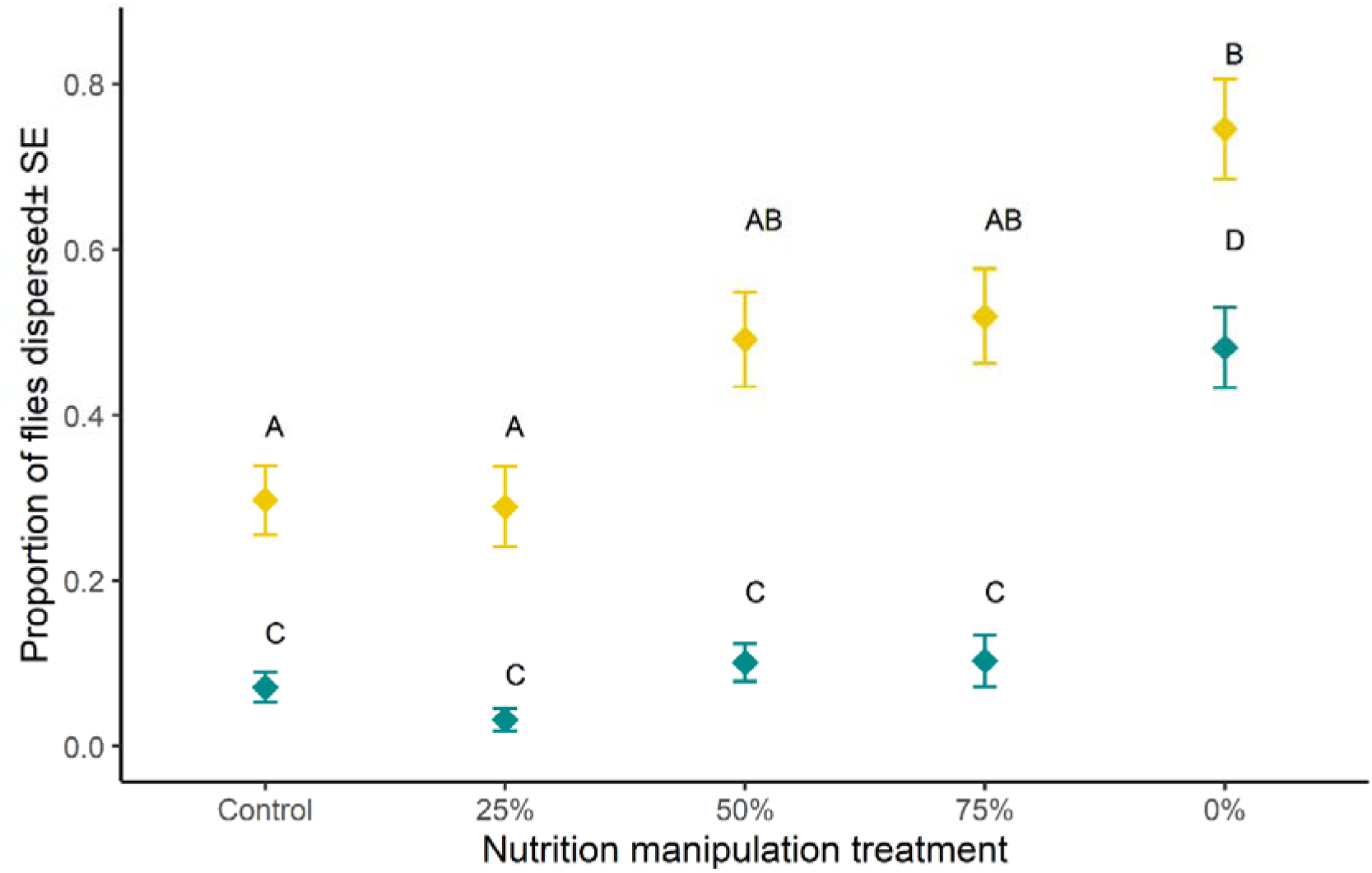
Results from Experiment 1.1. The effect of nutritional manipulation on dispersal tendency. Source patch conditions were varied by diluting all the nutritional components of standard food and thus creating - Control or, Standard food, 75%, 50%, 25% of standard food and 0% that is only non-nutritive agar media. Proportion of females (denoted with yellow colour) and males (denoted with grey colour), within a replicate two-patch setup, that dispersed from the source to sink at the end of 6 H is considered as a measure of dispersal (tendency) of that replicate and was used as the unit of analysis. The plots represent data combining all three blocks.

**Table 1:** Summary of the analyses of results from all experiments. The results were analysed using general linear mixed-effect models and binomial family distribution. Statistically significant p-values are mentioned in bold font style.

| Contrast | | $\chi^2$ | DF | p-value |
| --- | --- | --- | --- | --- |
| **Experiment 1.1: Effect of diet and nutrition on dispersal tendency** |  |  |  |  |
| Diet treatment |  | 39.33 | 4 | **<0.01** |
| Sex |  | 158.475 | 1 | **<0.01** |
| Diet treatment × Sex | | 65.13 | 4 | **<0.01** |
| **Experiment 1.2: The effect of protein and carbohydrate manipulation on dispersal tendency of female** |  |  |  |  |
| Diet treatment |  | 12.62 | 4 | **<0.01** |
| **Experiment 2.1: The effect of adult density on dispersal tendency** |  |  |  |  |
| Density treatment |  | 6.48 | 2 | **0.03** |
| Sex |  | 278.131 | 1 | **<0.01** |
| Density treatment × Sex | | 20.64 | 2 | **<0.01** |
| **Experiment 2.2: The effect of adult sex ratio on dispersal tendency** |  |  |  |  |
| Female dispersal | Sex ratio treatment | 8.07 | 3 | **0.04** |
| Male dispersal | Sex ratio treatment | 30.16 | 3 | **<0.01** |

For the second assay (Experiment 1.2), we found a significant effect of Treatment. Pairwise comparison showed that, from the ‘minimum protein’ level, flies dispersed significantly more compared to other levels of macronutrient treatment (except 25% protein level). In fact, 47% more females dispersed from minimum protein level (mean ±se: 0.65±0.04) compared to the control (mean ±se: 0.44±0.04) (Figure 3, Table 1).

**Figure 3:**
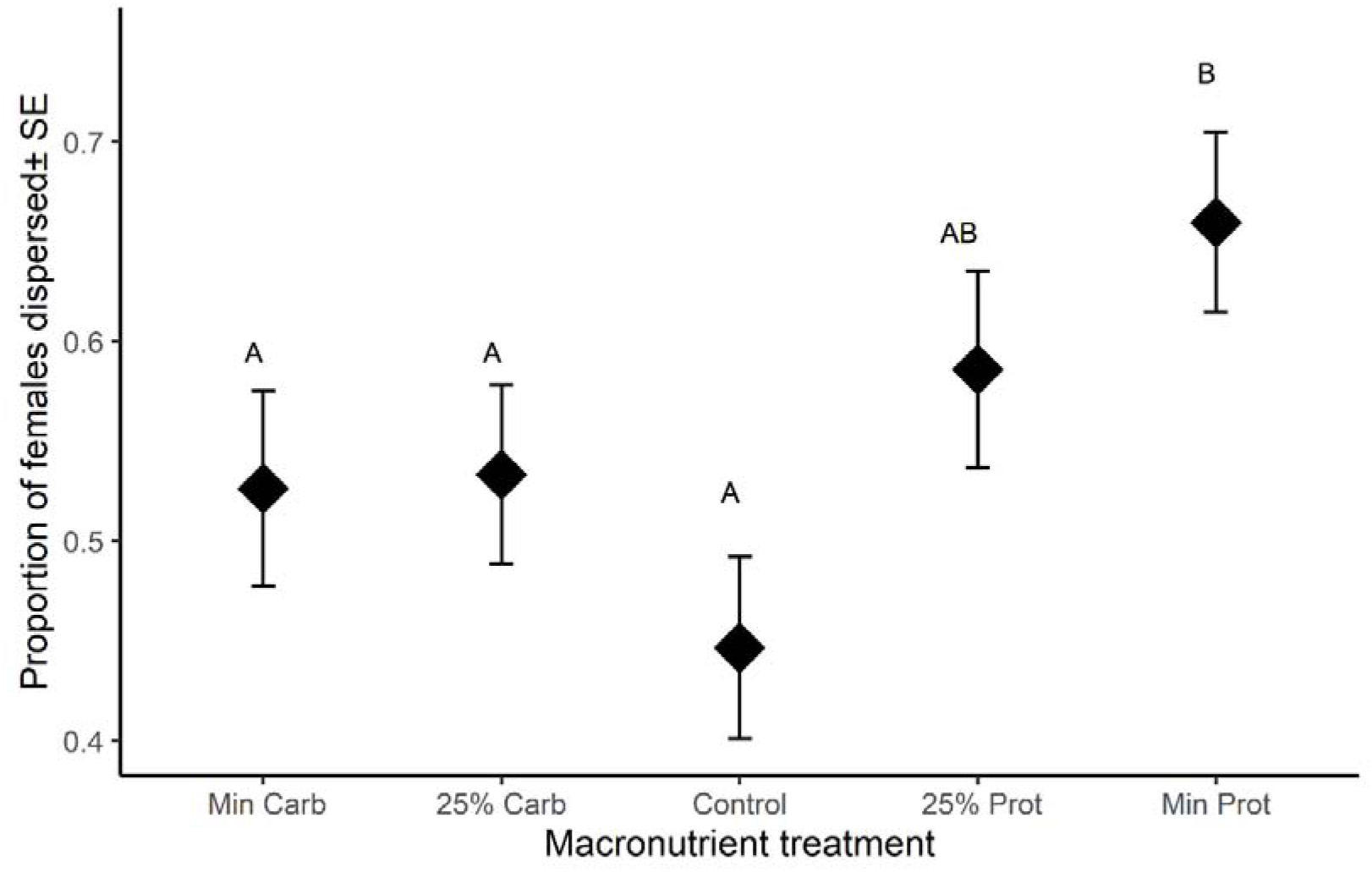
Results from Experiment 1.2. Effect of macronutrient manipulation treatment on dispersal tendency. The effect of protein and carbohydrate manipulation on dispersal tendency of females. Source patches treatment were of following types-Control or Standard food, 25% protein of Control, Minimum protein, 25% carbohydrate of Control, Minimum carbohydrate.

### Experiment 2

In the experiment 2.1, we found significant effects of Treatment, Sex and Treatment × Sex (Table 1). Females’ dispersal tendency did not differ across the levels of density treatment however, from the high density condition (mean ±se: 0.22±0.02) males dispersal tendency increased to 100% compared to standard or control density (mean ±se: 0.11±0.02) (Figure 4).

**Figure 4:**
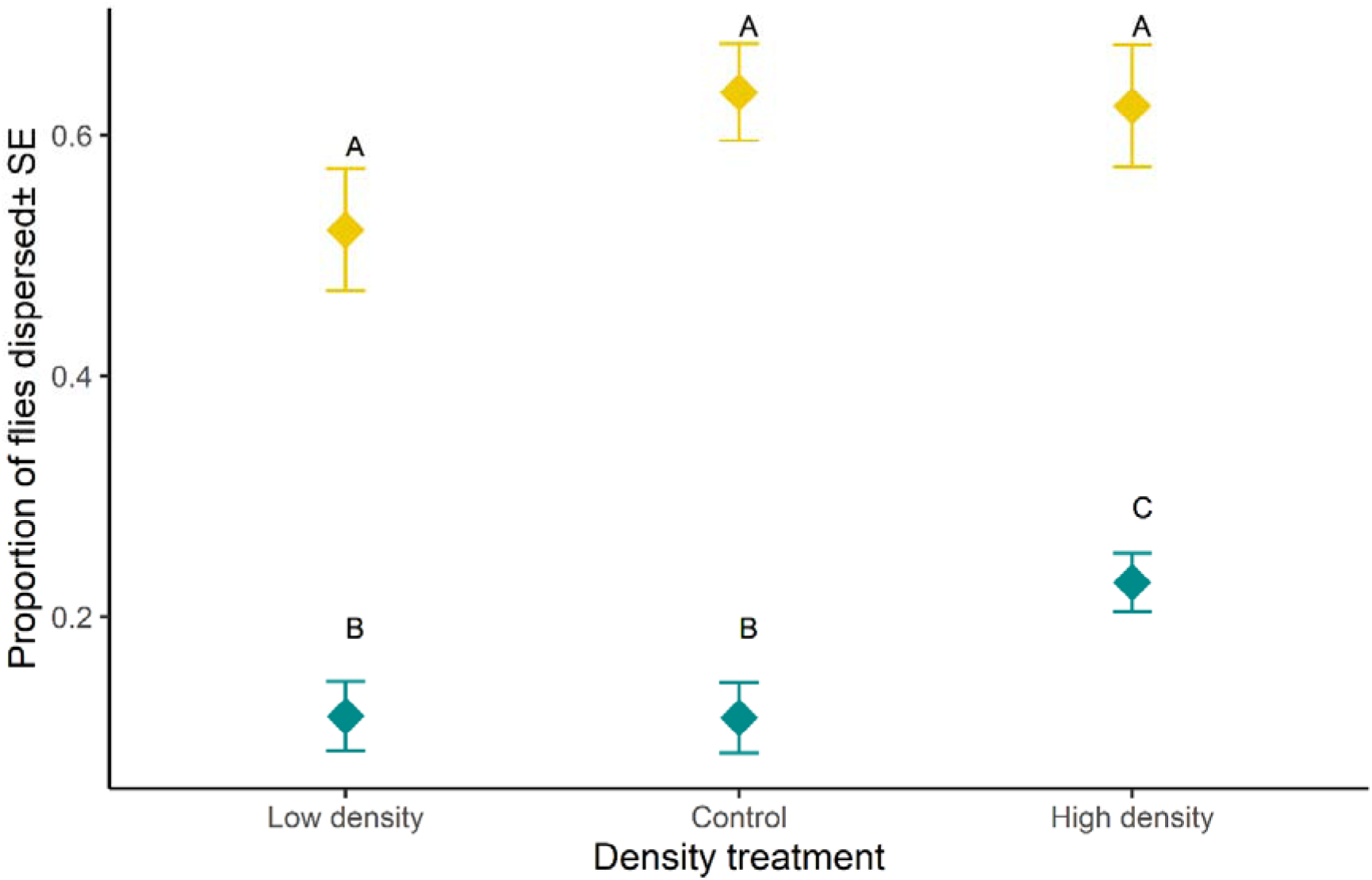
Results from Experiment 2.1. Effect of density treatment on dispersal tendency. The effect of adult density on dispersal tendency. Proportion of females (denoted with yellow colour) and males (denoted with grey colour), within a replicate two-patch setup, that dispersed from the source to sink at the end of 6 H is considered as a measure of dispersal (tendency) of that replicate and was used as the unit of analysis. The plots represent data combining all three blocks.

In experiment 2.2, we found a significant effect of sex ratio treatment for both female (Table 1, Figure 5) and male (Table 1, Figure 5) dispersal. Post-hoc multiple comparison indicates that females disperses significantly more from the male biased group than from all-female group (mean ±se: MB: 0.61 ±0.04, AF: 0.46±0.03) and with non-significant difference with other two levels i.e., FB and ES (Figure 5). In terms of males, dispersal from the all-male (AM) group was found to be significantly higher from the rest of three levels of sex ratio treatment (Figure 5). Males dispersal shoots up to 137.5 % in complete absence of females compared to the controlled sex-ratio condition (mean ±se: AM: 0.19±0.02, ES: 0.08 ±0.02).

**Figure 5:**
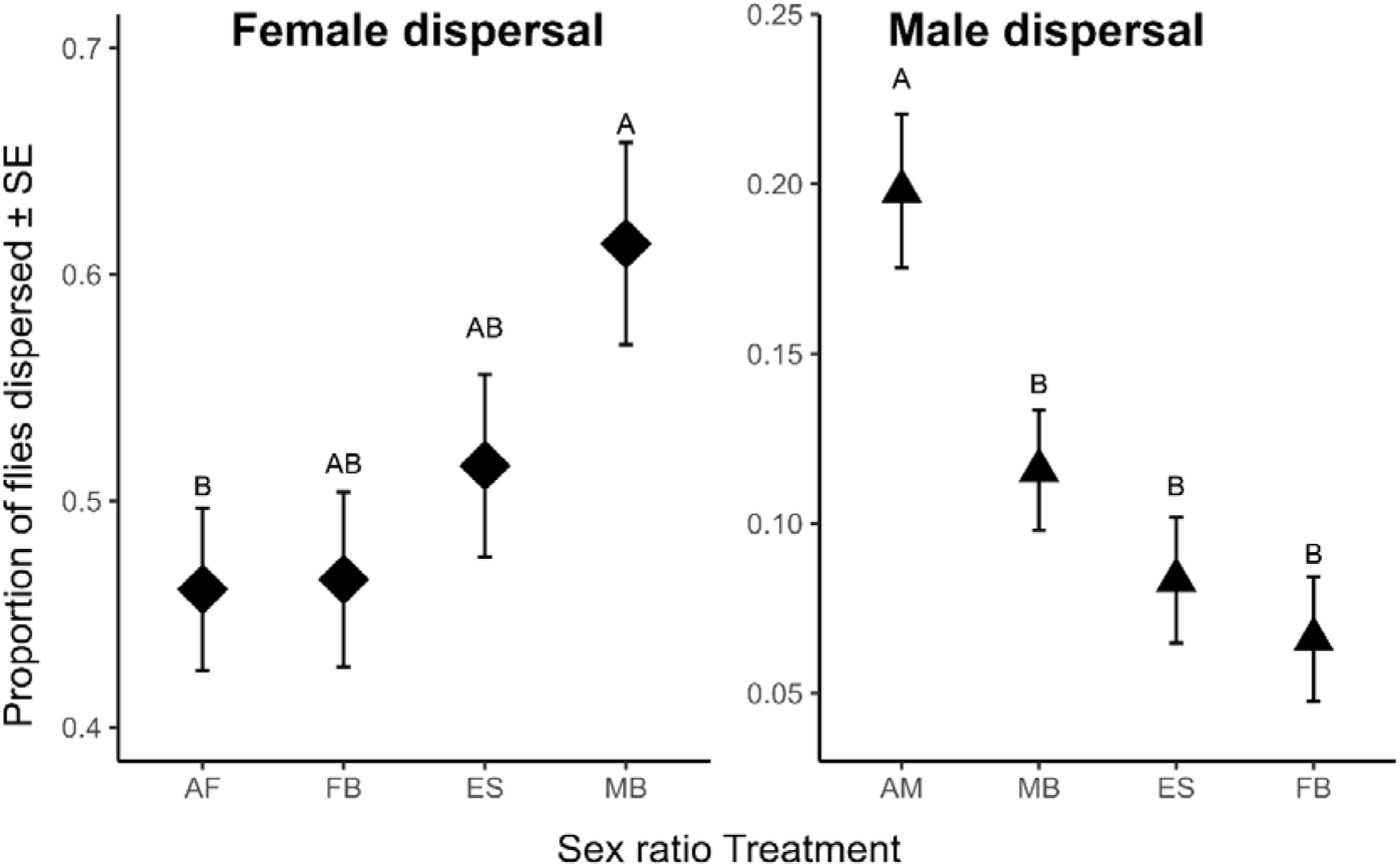
Results from Experiment 2.2. The effect of adult sex ratio on dispersal tendency. Varying sex ratio treatments were Standard or equal sex ratio (referred as ES), Male Biased (referred as MB), Female biased (referred as FB), All female group (referred as AF), All male group (referred as AM). Proportion of females and males within a replicate two-patch setup, that dispersed from the source to sink at the end of 6 H is considered as a measure of dispersal (tendency) of that replicate and was used as the unit of analysis. The plots represent data combining all three blocks.

## Discussion

With ample data on spontaneous dispersal using our setup, we have established an experimental paradigm to investigate Evolutionary Ecology of inter-patch dispersal and spatial structure of a population. Female-biased dispersal was found to be a consistent observation, across different experimental treatment. As theoretically predicted female dispersal tendency was found to be respond to diet treatment, especially to protein content, whereas such a trend was not seen in males unless they were exposed to starvation. While both density and sex ratio affected dispersal tendency, the effects were sex specific. High adult density increased dispersal in males, but not in females. Females dispersal was significantly higher under a male-biased sex ratio, whereas male dispersal was significantly higher in absence of females but otherwise unaffected by sex ratio.

Female fruit flies are reliant on transient food sources for nutrition, and also as oviposition substrate (Reaume & Sokolowski, 2006). Potentially, males can defend a territory and secure mating (Jonsson et al., 2011). Since female fitness critically depend on access to good quality food patches, female biased dispersal is not unexpected. Further, ephemeral food patches are expected to deplete relatively quickly in terms of source of nutrition as well as potential oviposition site. However, literature also suggest detrimental effect of higher locomotor activity on female fitness even in laboratory microcosm (Long & Rice, 2007). It is curious that such female biased dispersal pattern was not reported consistently in previous studies. These previous experiments using two-patch setup represent dispersal under a particularly extreme condition, viz., extreme starvation and desiccation. How much of these results can be extended to spontaneous dispersal, which is often exploratory in nature, is not clear. Nonetheless, in spite of the dominant effect of starvation and/or desiccation, at least some of these studies showed density dependent sex biased dispersal (Wang et al., 2014; Mishra et al., 2022).

There can be at least two potential alternative explanation of the observed pattern of female biased dispersal in our assays. First, females in *D. melanogaster* are, on an average, larger than males (Horabin, 2005), and are hence expected to have higher amount of stored resources in their body (Gerber et al., 2022). Body size dependent dispersal rate is not unheard of (Bonte et al., 2012, see Anderson and Fahimipour, 2021 for a relatively recent exposition relevant to stability of metacommunity). Hence, in principle, it is possible that sex difference in dispersal behaviour observed in our experiments only reflects sexual size dimorphism. However, this is very unlikely. First, sexual size dimorphism is a common attribute of all *D. melanogaster* systems – and yet, this is only the first time female biased dispersal propensity is being reported explicitly. In addition, we showed that there is consistent variation in body size (see Figure S2) without a corresponding change in dispersal propensity (Halder et al., 2024). Secondly, since our measure of dispersal propensity was reliant on actual movement of individuals, if our two-patch dispersal setup allowed more female movement, it can result into the observed pattern of results. However, if our dispersal setup allows females to reach sink freely, it is unlikely that the same setup restricted movement of males which are generally significantly smaller in size.

Since dispersal is a physiologically costly behaviour (Bonte et al. 2012), expression of it can be expected to condition dependent. Therefore, nutritive status, juvenile growth condition etc., are likely to affect dispersal propensity. Fruit fly females are known to rely heavily on foraging, especially on yeast, during the adult stage (Chippindale et al., 1993, Anagnostou et al., 2010). Therefore, it is possible that spontaneous dispersal to have evolved as an adaptive strategy allowing the females to explore other food patches with a finite probability of discovering a patch with better reward. Interestingly, changing the carbohydrate (jaggary in our fly food) content, which effectively alter the caloric value of the food, did not affect female dispersal, but alteration of protein (yeast in our fly food) content did. Since, female fecundity is mainly limited by access to protein diet, this is suggestive of adaptive female dispersal behaviour that facilitate the attainment of maximum fecundity.

It is important to note that the observed dispersal pattern could not have been due to nutritional deprivation as 6 hours of exposure to impoverished diet, as was done during the assay, is unlikely to result in nutrient deprivation as such. Therefore, females’ response could be a response based on the perception of food quality. It remains to be seen if such dispersal can be triggered just by the cue associated with such nutritionally impoverished food.

Since food is also the oviposition site for females, cost of not moving from a poor quality food patch can be quite high for them. Results from our pilot studies suggest that poor larval diet, especially 40% dilution of nutritional value, can lead to a reduction offspring survival by 6.25 % (Mean ± SE: Control: 0.85± 0.00; 40%: 0.80±0.01; see supplementary information, Table S6, Figure S3), and in addition can prolong egg-to-adult development time (Table S4, Figure S1) and reduce adult body weight at eclosion (Table S5, Figure S2). Therefore, practically all fitness components of the offspring are likely to be affected if a female chooses to deposit eggs on a nutritionally depleted food. It is known that females can disperse in search of potential oviposition site (Lihoreau et al., 2016; Camus et al., 2018). Fruit fly females are known to show oviposition site preference (Fanara et al., 2023). Nonetheless, the idea of female dispersal in search of suitable oviposition site needs further investigations.

Males in this system are known to show little foraging behaviour (Long & Rice., 2007). Hence, nutrient dependence of male fitness and associated behavioural traits is expected to stem from juvenile developmental nutrition (Zajitschek et al., 2016). Previously, we have shown that larval dietary manipulation treatment had a small but significant effect on adult spontaneous dispersal in males (Halder et al., 2024). It appears that the observed pattern of male dispersal tendency is best explained by intra-sexual selection (i.e., male-male competition) rather than condition dependence.

Density and sex ratio, both can trigger dispersal by introducing variation in intraspecific competition and intersexual interactions (Kokko & Rankin 2006). It is important to note that these two aspects of a population are not independent. While density regulates the extent of intraspecific competition regardless of sex (Aronsen et al., 2013), sex ratio can affect a specific type of intraspecific competition – viz., male-male competition, especially at male biased vs. female biased operational sex ratio (Trochet et al., 2013, Baines et al., 2017). High male-male competition results in expression of competitive male traits, which have been often found to result in detrimental side-effect in females, resulting in intersexual conflict (Rice et al., 2006; Wolfner, 2009; Nandy et al., 2013). Our results on density effect on dispersal suggested a sex specific effect. While female dispersal was found to be largely density independent, males showed higher dispersal at high density. This discord between sexes appears to fit with the idea that in our laboratory setup females rarely compete (for food) as the conditions are generally *ad-lib*, while males are always under competition for mating opportunity (Prasad and Joshi 2003, Rice et al. 2006). Under high density and in male-biased sex ratio, male-male competition is expected to peak. Hence, we argue that higher male dispersal under high density was likely due to increased male-male competition. However, male-biased sex ratio was not sufficient to generate higher male dispersal, which appeared only under complete lack of mates (i.e., under all-male condition). Such phenomenon has been previously referred to as “mate-finding dispersal” (Mishra et al. 2020). It will be interesting to know whether male dispersal is based on territoriality such that competitively weaker males disperse more often.

Trochet et al. (2013) did not find evidence of sex-biased dispersal in cabbage butterfly (*Pieris brassicae*) by varying adult sex ratio, but suggested that male harassment (a by-product of male-male competition) as a potential driving factor for the observed dispersal in females. Detrimental effect of male inflicted harm to females in terms of reduced fecundity, survival, and stress tolerance is well known (Chapman et al., 1995; Rice et al., 2006; Wolfner, 2009; Nandy et al., 2013). In *D. melanogaster,* females are known to resist mating by kicking, flicking or extrusion of genitalia (Connolly & Cook, 1973). In a spatially structured habitat, one simple mate avoidance strategy could be to move away from a patch to a neighbouring patch – either avoiding male-prevalent patch or seeking refuge (Yun et al., 2017, Halder et al., 2024). We found females to disperse more from a male-biased source patch compared to all female group. This fits well with the mate-avoidance female dispersal theory. Interestingly, in a spatially structured microcosm habitat with inter-patch dispersal, sexual-conflict does not necessarily change rather inflicts novel fitness cost due to energetically demanding movement (Halder et al., 2024).

## Acknowledgements

We are thankful to the Indian Institute of Science Education and Research Berhampur for a generous intramural grant that provided the financial support for this investigation. SH is thankful to the University Grant Commission, Govt. of India for financial support in the form of Junior and Senior Research Fellowship. We thank Purbasha Dasgupta, Tanya Verma, and Rabi Sankar Pal for their help in the lab during the assay.

## References

Anagnostou C, Dorsch M, Rohlfs M. (2010) Influence of dietary yeasts on Drosophila melanogaster lifelJhistory traits. Entomologia Experimentalis et Applicata. 136:1–1.

Anderson KE, Fahimipour AK. (2021) Body size dependent dispersal influences stability in heterogeneous metacommunities. Scientific reports. 11:17410.

Aronsen T, Berglund A, Mobley KB, Ratikainen II, Rosenqvist G. (2013) Sex ratio and density affect sexual selection in a sex-role reversed fish. Evolution. 67:3243–57.

Baines CB, Ferzoco IM, McCauley SJ. (2017) Sex-biased dispersal is independent of sex ratio in a semiaquatic insect. Behavioral Ecology and Sociobiology. 71:1–7.

Bateman, A. J. (1948) Intra-sexual selection in Drosophila. Heredity. 2:349–368.

Berglund A. (1995) Many mates make male pipefish choosy. Behaviour. 132:213–8.

Bitume EV, Bonte D, Ronce O. (2013) Density and genetic relatedness increase dispersal distance in a subsocial organism. Ecol Lett. 16:430–437.

Bonte D, Van Dyck H, Bullock JM, Coulon A, Delgado M, Gibbs M, Travis JM. (2012) Costs of dispersal. Biological reviews. 87: 290–312.

Bowler DE, Benton TG. (2005) Causes and consequences of animal dispersal strategies: Relating individual behaviour to spatial dynamics. Biol Rev Camb Philos Soc. 80:205–225.

Camus MF, Huang CC, Reuter M, Fowler K. (2018) Dietary choices are influenced by genotype, mating status, and sex in Drosophila melanogaster. Ecology and Evolution. 8:5385–93.

Chapman T, Liddle LF, Kalb JM, Wolfner MF, Partridge L. (1995) Cost of mating in Drosophila melanogaster females is mediated by male accessory gland products. Nature. 373: 241–244.

Chippindale AK, Leroi AM, Kim SB, Rose MR. (1993) Phenotypic plasticity and selection in Drosophila lifelJhistory evolution. I. Nutrition and the cost of reproduction. Journal of Evolutionary Biology. 6:171–93.

Clarke AL, Sæther BE, Røskaft E. (1997) Sex biases in avian dispersal: a reappraisal. Oikos. 429–438.

Clutton-Brock TH, Russell AF, Sharpe LL, Young AJ, Balmforth Z, McIlrath GM. (2002) Evolution and development of sex differences in cooperative behavior in meerkats. Science. 297:253–6.

Clobert J, Le Galliard JF, Cote J, Meylan S, Massot M. (2009) Informed dispersal, heterogeneity in animal dispersal syndromes and the dynamics of spatially structured populations. Ecology letters. 12:197–209.

Clobert J, editor. (2012) Dispersal ecology and evolution. Oxford University Press.

Connolly K, Cook R. (1973) Rejection responses by female Drosophila melanogaster: their ontogeny, causality and effects upon the behaviour of the courting male. Behaviour. 1:142–66.

Cotter SC, Littlefair JE, Grantham PJ, Kilner RM. (2013) A direct physiological tradelJoff between personal and social immunity. Journal of Animal Ecology. 82:846–53.

Coyne JA, Boussy IA, Prout T, Bryant SH, Jones JS, Moore JA. (1982) Long-distance migration of Drosophila. The American Naturalist. 119:589–95.

Coyne JA, Bryant SH, Turelli M. (1987) Long-distance migration of Drosophila. 2. Presence in desolate sites and dispersal near a desert oasis. The American Naturalist. 129:847–61.

Dasgupta P, Halder S, Dari D, Nabeel P, Vajja SS, Nandy B. (2022) Evolution of a novel female reproductive strategy in Drosophila melanogaster populations subjected to longlJterm protein restriction. Evolution. 76:1836–48.

Dobson FS. (1982) Competition for mates and predominant juvenile male dispersal in mammals. Anim Behav. 30:1183–92.

Dobzhansky T, Wright S. (1943) Genetics of natural populations. X. Dispersion rates in Drosophila pseudoobscura. Genetics. 28:304.

Dukas R. (2020) Natural history of social and sexual behavior in fruit flies. Scientific reports. 10:21932.

Fanara JJ, Beti MI, Gandini L, Hasson E. (2023) Oviposition behaviour in Drosophila melanogaster: Genetic and behavioural decoupling between oviposition acceptance and preference for natural fruits. Journal of Evolutionary Biology. 36:251–63.

Friedenberg NA. (2003) Experimental evolution of dispersal in spatiotemporally variable microcosms. Ecology Letters. 6:953–9.

Fronhofer EA, Legrand D, Altermatt F. (2018) Bottom-up and top-down control of dispersal across major organismal groups. Nat Ecol Evol. 2:1859–1863.

Gerber R, Cabon L, Piscart C, Roussel JM, Bergerot B. (2022) Body stores of emergent aquatic insects are associated with body size, sex, swarming behaviour and dispersal strategies. Freshwater Biology. 67:2161–75.

Greenwood PJ. (1980) Mating systems, philopatry and dispersal in birds and mammals. Anim Behav. 28:1140–62.

Halder S, Kar S, Sethi S, Tewari S, Verma T, Nandy B. (2022) Spatial structure does not reduce population level interlocus sexual conflict but imposes additional sex-specific costs.

Horabin JI. (2005) Splitting the Hedgehog signal: sex and patterning in Drosophila. 4801–4810

Iliadi KG., Iliadi NN, Rashkovetsky EL, Girin SV, Nevo E, Korol AB. (2002) Sexual differences for emigration behavior in natural populations of Drosophila melanogaster. Behavior genetics. 32: 173–180.

Jirotkul M. (1999) Population density influences male–male competition in guppies. Animal Behaviour. 58:1169–75.

Jonsson T & Kravitz EA. (2011) Heinrich R. Sound production during agonistic behavior of male Drosophila melanogaster. Fly. 5:29–38.

Jungwirth A, Zöttl M, Bonfils D, Josi D, Frommen JG, Taborsky M. (2023) Philopatry yields higher fitness than dispersal in a cooperative breeder with sex-specific life history trajectories. Science Advances. 9:eadd2146.

Li XY, Kokko H. (2019) Intersexual resource competition and the evolution of sex-biased dispersal. Front Ecol Evol. 7:1–9.

Li XY, Kokko H. (2019) Sex-biased dispersal: a review of the theory. Biol Rev. 94:721–736.

Lihoreau M, Poissonnier LA, Isabel G, Dussutour A. (2016) Drosophila females trade off good nutrition with high-quality oviposition sites when choosing foods. J Exp Biol. 219:2514–2524.

Long TA & Rice WR. (2007) Adult locomotory activity mediates intralocus sexual conflict in a laboratory-adapted population of Drosophila melanogaster. Proceedings of the Royal Society B: Biological Sciences. 274:3105–12.

Mabry KE, Shelley EL, Davis KE, Blumstein DT, Van Vuren DH. (2013) Social mating system and sex-biased dispersal in mammals and birds: a phylogenetic analysis. PloS one. 8:e57980.

Mishra A, Tung S, Shreenidhi PM, AamirSadiq M, Shree Sruti VR, Chakraborty PP, Dey S. (2018) Sex differences in dispersal syndrome are modulated by environment and evolution. Philosophical Transactions of the Royal Society B: Biological Sciences. 373: 20170428.

Mishra A, Tung S, Sruti VS, Sadiq MA, Srivathsa S, Dey S. (2018) PrelJdispersal context and presence of opposite sex modulate density dependence and sex bias of dispersal. Oikos. 127: 1596–1604.

Mishra A, Tung S, Shree Sruti VR, Srivathsa S, Dey S. (2020) MatelJfinding dispersal reduces local mate limitation and sex bias in dispersal. Journal of Animal Ecology. 89: 2089–2098.

Mishra A, Tung S, Sruti VS, Shreenidhi PM, Dey S. (2022) Desiccation stress acts as cause as well as cost of dispersal in Drosophila melanogaster. The American Naturalist. 199: E111–E123.

Nandy B, Gupta V, Sen S, Udaykumar N, Samant MA, Ali SZ, Prasad NG. (2013) Evolution of mate-harm, longevity and behaviour in male fruit flies subjected to different levels of interlocus conflict. BMC evolutionary biology. 13: 1–16.

Partridge L, Hoffmann A, Jones JS. (1987) Male size and mating success in Drosophila melanogaster and D. pseudoobscura under field conditions. Animal Behaviour. 35:468–76.

Partridge L, Fowler K. (1990) Non-mating costs of exposure to males in female Drosophila melanogaster. Journal of Insect Physiology. 36:419–25.

Perrin N, Mazalov V. (2000) Local competition, inbreeding, and the evolution of sex-biased dispersal. Amer Naturalist. 155:116–27

Prasad NG, Joshi A. (2003) What have two decades of laboratory life-history evolution studies on Drosophila melanogaster taught us? Journal of genetics. 82:45–76.

Rapkin J, Jensen K, Archer CR, House CM, Sakaluk SK, Castillo ED, Hunt J. (2018) The geometry of nutrient space–based life-history trade-offs: sex-specific effects of macronutrient intake on the trade-off between encapsulation ability and reproductive effort in decorated crickets. The American Naturalist, 191: 452–474.

Reaume CJ, Sokolowski MB. (2006) The nature of Drosophila melanogaster. Current biology. 16:R623–8.

Reyes E., Cunliffe F., & M’Gonigle, L K. (2023) Evolutionary dynamics of dispersal and local adaptation in multi-resource landscapes. Theoretical Population Biology, 153: 102–110.

Rice WR, Stewart AD, Morrow EH, Linder JE, Orteiza N, Byrne PG. (2006) Assessing sexual conflict in the Drosophila melanogaster laboratory model system. Philosophical Transactions of the Royal Society B: Biological Sciences. 361: 287–299.

Ronce O. (2007) How does it feel to be like a rolling stone? Ten questions about dispersal evolution. Annu. Rev. Ecol. Evol. Syst.. 231–53.

Stevens VM, Trochet A, Van Dyck H, Clobert J. (2012) Baguette M. How is dispersal integrated in life histories: a quantitative analysis using butterflies. Ecology letters. 1:74–86.

Tompkins L, Gross AC, Hall JC, Gailey DA, Siegel RW. (1982) The role of female movement in the sexual behavior of Drosophila melanogaster. Behavior genetics. 12:295–307.

Travis JMJ, Delgado M, Bocedi G. (2013) Dispersal and species’ responses to climate change. Oikos. 122:1532–1540.

Trochet A, Legrand D, Larranaga N, Ducatez S, Calvez O, Cote J, Clobert J, Baguette M. (2013) Population sex ratio and dispersal in experimental, twolJpatch metapopulations of butterflies. Journal of Animal Ecology. 82:946–55.

Trochet A, Courtois EA, Stevens VM, Baguette M, Chaine A, Schmeller DS, Wiens JJ. (2016) Evolution of sex-biased dispersal. The Quarterly Review of Biology. 91: 297–320

Tung S, Mishra A, Gogna N, Aamir Sadiq M, Shreenidhi PM, Shree Sruti VR, Dorai K, Dey S. (2018) Evolution of dispersal syndrome and its corresponding metabolomic changes. Evolution. 72:1890–903.

Tung S, Mishra A, Shreenidhi PM, Sadiq MA, Joshi S, Sruti VS, Dey S. (2018) Simultaneous evolution of multiple dispersal components and kernel. Oikos. 127:34–44.

Van Hooft P, Keet DF, Brebner DK, Bastos AD. (2018) Genetic insights into dispersal distance and disperser fitness of African lions (Panthera leo) from the latitudinal extremes of the Kruger National Park, South Africa. BMC genetics. 19:1–6.

Venkateswaran V, Shrivastava A, Kumble AL, Borges RM. (2017) Life-history strategy, resource dispersion and phylogenetic associations shape dispersal of a fig wasp community. Movement Ecology. 5:1–1.

Wang S-P, Guo W-Y, Muhammad SA, Chen R-R, Mu L-L, Li G-Q. (2014) Mating experience and food deprivation modulate odor preference and dispersal in Drosophila melanogaster males. J Insect Sci. 14:1–14.

Woods JK, Kowalski S, Rogina B. (2014) Determination of the spontaneous locomotor activity in Drosophila melanogaster. JoVE (Journal of Visualized Experiments). 10:e51449.

Wolff JO, Plissner JH. (1998) Sex biases in avian natal dispersal: an extension of the mammalian model. Oikos. 83:327–30.

Wolfner MF. (2009) Battle and ballet: molecular interactions between the sexes in Drosophila. Journal of Heredity.100: 399–410.

Yun L, Chen PJ, Singh A, Agrawal AF, Rundle HD. (2017) The physical environment mediates male harm and its effect on selection in females. Proceedings of the Royal Society B: Biological Sciences. 284: 20170424.

Zajitschek F, Zajitschek SR, Canton C, Georgolopoulos G, Friberg U, Maklakov AA. (2016) Evolution under dietary restriction increases male reproductive performance without survival cost. Proceedings of the Royal Society B: Biological Sciences. 283:20152726.

Zajitschek F, Georgolopoulos G, Vourlou A, Ericsson M, Zajitschek SR, Friberg U, Maklakov AA. (2019) Evolution under dietary restriction decouples survival from fecundity in Drosophila melanogaster females. The Journals of Gerontology: Series A. 74:1542–8.

