## Supplementary information file for "Nutrition and density dependence of spontaneous female-biased dispersal in *Drosophila melanogaster*"

**Table S1: Food composition and recipe for Experiment 1.** Control or Standard food, 75%, 50%, 25% of standard food and 0% that is only non-nutritive agar media

| **Ingredients** | **Control** | **75% Diluted** | **50% Diluted** | **25% Diluted** | **0%** |
| --- | --- | --- | --- | --- | --- |
| **Banana (g)** | 205 | 153.75 | 102.5 | 51.25 | 0 |
| **Barley (g)** | 25 | 18.75 | 12.5 | 6.25 | 0 |
| **Jaggery (g)** | 35 | 26.25 | 17.5 | 8.75 | 0 |
| **Yeast (g)** | 36 | 27 | 18 | 9 | 0 |
| **Agar-Agar (g)** | 12.4 | 12.4 | 12.4 | 12.4 | 12.4 |
| **Water (mL) [to be mixed with Agar-Agar]** | 1000 | 1000 | 1000 | 1000 | 1000 |
| **Ethanol (mL) [to be mixed with Yeast]** | 22 | 22 | 22 | 22 | 22 |
| **Water (mL)** | 180 | 180 | 180 | 180 | 180 |
| **p-Hydroxy Methyl benzoate (g)** | 2.4 | 2.4 | 2.4 | 2.4 | 2.4 |
| **Ethanol (mL) [to be mixed with Benzoate]** | 23 | 23 | 23 | 23 | 23 |

**Table S2: Food composition and recipe for Experiment 2.** Control or Standard food, 25% protein of Control, Minimum protein, 25% carbohydrate of Control, Minimum carbohydrate.

| **Ingredients** | **Control** | **25% Jaggery** | **25% Yeast** | **Min Carbohydrate** | **Min Protein** |
| --- | --- | --- | --- | --- | --- |
| **Banana (g)** | 205 | 205 | 205 | 205 | 205 |
| **Barley (g)** | 25 | 25 | 25 | 25 | 25 |
| **Jaggery (g)** | 35 | 8.75 | 35 | 0 | 35 |
| **Yeast (g)** | 36 | 36 | 9 | 36 | 0 |
| **Agar-Agar (g)** | 12.4 | 12.4 | 12.4 | 12.4 | 12.4 |
| **Water (mL) [to be mixed with Agar-Agar]** | 1000 | 1000 | 1000 | 1000 | 1000 |
| **Ethanol (mL) [to be mixed with Yeast]** | 22 | 22 | 22 | 22 | 22 |
| **Water (mL)** | 180 | 180 | 180 | 180 | 180 |
| **p-Hydroxy Methyl benzoate (g)** | 2.4 | 2.4 | 2.4 | 2.4 | 2.4 |
| **Ethanol (mL) [to be mixed with Benzoate]** | 23 | 23 | 23 | 23 | 23 |

Effect of larval diet on pre-adult development time, juvenile survivorship and dry bodyweight

Pre-adult (egg-to-adult) development time, juvenile (egg-to-adult) survivorship and dry weight at eclosion were measured for flies developed in Control, 20% and 40% larval diet (see table 3.9). Eggs were collected from the baseline population cages in exact numbers. For egg-collection, two small food plates (60mm diameter) were provided in a cage, and were left undisturbed for one hour. Following that those two plates were replaced by fresh two small food plates and again left undisturbed for one hour to allow oviposition. For the experiments, eggs were collected from the second set of plates. Such small oviposition window ensured the developmental synchrony of the eggs collected for the assay. Eggs were randomly collected and carefully counted on small piece of 1% Agar gel using a fine brush. Exact 70 eggs was taken and transferred to a fresh food vial having 8ml food which was either Control, 20% and 40% diluted food. Ten such vials were set up for each diet treatment. The entire experiment was performed in three independent statistical blocks using three randomly selected BL population (BL_3, 4, 5_). The vials were monitored carefully for the signs of eclosion. Beginning from the eclosion of the first individual, the eclosing adults were counted and sexed in 6-hour interval until all the pupae eclosed. From the time of setting up a vial, time to eclosion data was used to derive the average measure of egg-to-adult development time (i.e., pre-adult development time) for males and females in each vial. In addition, out of the 70 eggs initially introduced, the proportion of flies that finally eclosed from a vial was considered a measure of egg-to-adult survivorship (i.e., juvenile survivorship). For each block, 50 males and 50 females were collected for dry whole body weight measurement. They were collected within 6 hours of eclosion from the peak of eclosion, and were immediately frozen at -20°C. Flies were later dried at 60°C for 48 hr and weighed in groups of five using Shimadzu AUW220D to the nearest 0.01 mg. The mean body weight of the group of 5 was taken as a unit of analysis.

Data on egg-to-adult development time, body weight at eclosion and egg-to-adult survival was analysed using linear model (LM). Analysis was done with treatment (Larval diet), sex and block as fixed effect by using package lme4 and function lm in R (R Version 4.2.0). Analysis of pre-adult development time and body weight showed significant effect of treatment × block interaction. Therefore, we re-analysed development time and body weight for each block separately. Data of egg-to-adult development time for block 3 was not normally distributed and hence were analysed using generalized linear model (GLM), function glm with gamma family distribution. For survivorship analysis we did not find significant effect of treatment × block interaction. Hence, all the blocks were pooled together. Whenever significant effect of treatment or treatment × sex interaction was found, pairwise comparisons were done using the package emmeans.

Though the degree of response varied from population to population, significant effect of treatment on measured traits in all the blocks suggest that poor larval diet such as 40% dilution can adversely affect not only egg-to-adult survival (Mean ± SE: Control: 0.85± 0.00; 40%: 0.80±0.01; Table 3.13, Figure 3.10) but also egg-to-adult development time (Table 3.11, Figure 3.8) and body weight at eclosion (Table 3.12, Figure 3.9).

**Table S3: Food composition and recipe for larval diet.** Control (i.e., complete diet), and 20%, 40%, 60% dilution of the nutritional components of the standard food.

| **Ingredients** | **Control** | **20% diluted** | **40% diluted** | **60% diluted** |
| --- | --- | --- | --- | --- |
| **Banana (gm)** | 205 | 164 | 123 | 82 |
| **Barley (gm)** | 25 | 20 | 15 | 10 |
| **Jaggery (gm)** | 35 | 28 | 21 | 14 |
| **Yeast (gm)** | 36 | 28.8 | 21.6 | 14.4 |
| **Agar-agar (gm)** | 12.4 | 12.4 | 12.4 | 12.4 |
| **Water (ml)**  **[to mix with agar-agar]** | 1000 | 1000 | 1000 | 1000 |
| **Ethanol (ml)**  **[to mix with yeast]** | 22 | 22 | 22 | 22 |
| **Water (ml)** | 180 | 180 | 180 | 180 |
| **p-Hydroxy methyl benzoate** | 2.4 | 2.4 | 2.4 | 2.4 |
| **Ethanol (ml)**  **[to mix with benzoate]** | 23 | 23 | 23 | 23 |

**Table S4: Summary of the analyses of results from development time.** The results were analysed using linear model or general linear model where larval diet treatments (treatment) and sex was fitted as fixed factors. Each Block was analysed seperately. Statistically significant p-values are mentioned in bold font style.

| **Development time** | **Blocks** | **Contrast** | $\mathbf{Sum sq}$ | **DF** | **p-value** |
| --- | --- | --- | --- | --- | --- |
|  | Block 1 | Treatment | 14417.2 | 2 | **<0.01** |
|  |  | Sex | 5564.2 | 1 | **<0.01** |
|  |  | Treatment × Sex | 31.1 | 2 | 0.79 |
|  | Block 2 | Treatment | 306.33 | 2 | **<0.01** |
|  |  | Sex | 594.98 | 1 | **<0.01** |
|  |  | Treatment × Sex | 6.52 | 2 | 0.51 |
|  | Block 3 | Treatment | 267.316 | 2 | **<0.01** |
|  |  | Sex | 49.865 | 1 | **<0.01** |
|  |  | Treatment × Sex | 11.840 | 2 | **<0.01** |

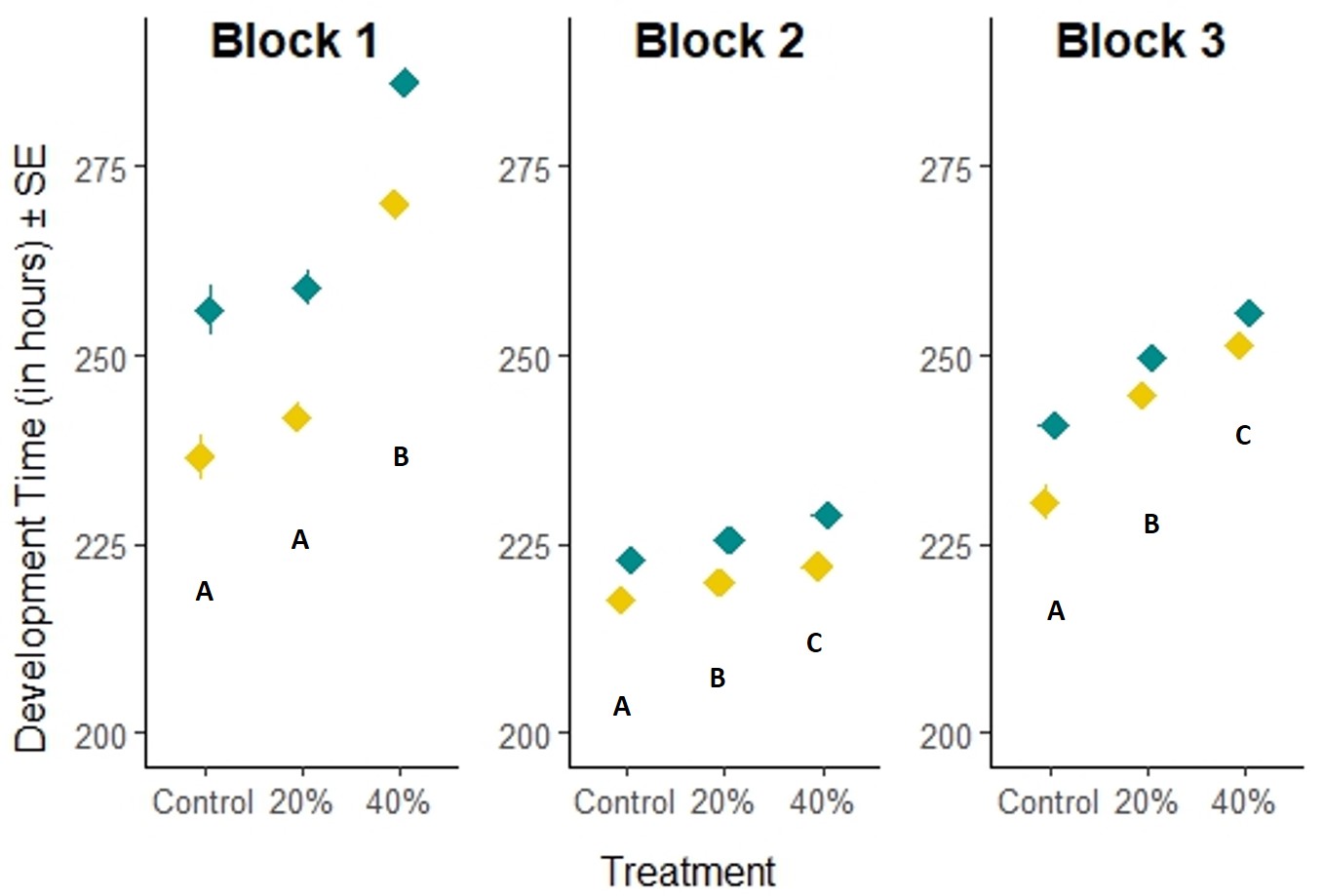

**Figure S1**: **Effect of larval diet treatment on Development time.** The effect of larval diet treatment on egg-to-adult development time. Treatments not sharing common alphabets are significantly different from each other. Development time of females are denoted with yellow color and males with cyan color.

**Table S5: Summary of the analyses of results from Body weight.** The results were analysed using linear model where diet treatments (treatment) and sex was fitted as fixed factors. Each Block was analysed seperately. Statistically significant p-values are mentioned in bold font style.

| **Body weight** | **Blocks** | **Contrast** | $\chi^{2}$ | **DF** | **p-value** |
| --- | --- | --- | --- | --- | --- |
|  | Block 1 | Treatment | 5510.8 | 2 | **<0.01** |
|  |  | Sex | 15690.7 | 1 | **<0.01** |
|  |  | Treatment × Sex | 226.6 | 2 | 0.26 |
|  | Block 2 | Treatment | 3920 | 2 | **<0.01** |
|  |  | Sex | 228597 | 1 | **<0.01** |
|  |  | Treatment × Sex | 365 | 2 | 0.18 |
|  | Block 3 | Treatment | 5097.6 | 2 | **<0.01** |
|  |  | Sex | 17094.6 | 1 | **<0.01** |
|  |  | Treatment × Sex | 159.6 | 2 | 0.38 |

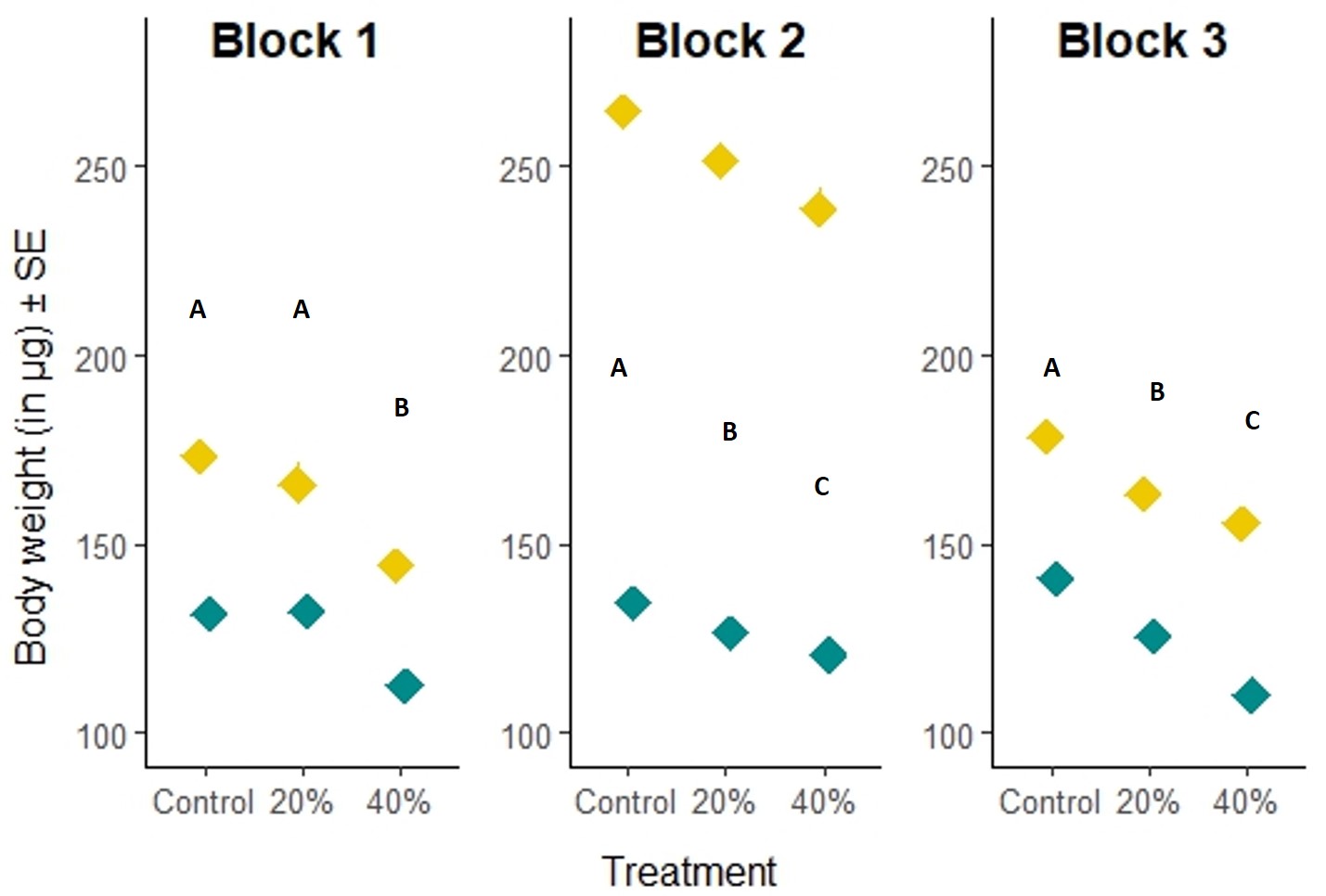

**Figure S2**: **Effect of larval diet treatment on Body weight.** The effect of adult density on body weight at eclosion. Body weight of females are denoted with yellow color and males with cyan color. Treatments not sharing common alphabets are significantly different.

**Table S6: Summary of the analyses of results from survivorship.** The results were analysed using linear models where diet treatments (treatment) was fitted as fixed factors. Statistically significant p-values are mentioned in bold font style.

| **Survivorship** | **Contrast** | **SumSq** | **DF** | **p-value** |
| --- | --- | --- | --- | --- |
|  | Treatment | 0.039 | 2 | **0.01** |

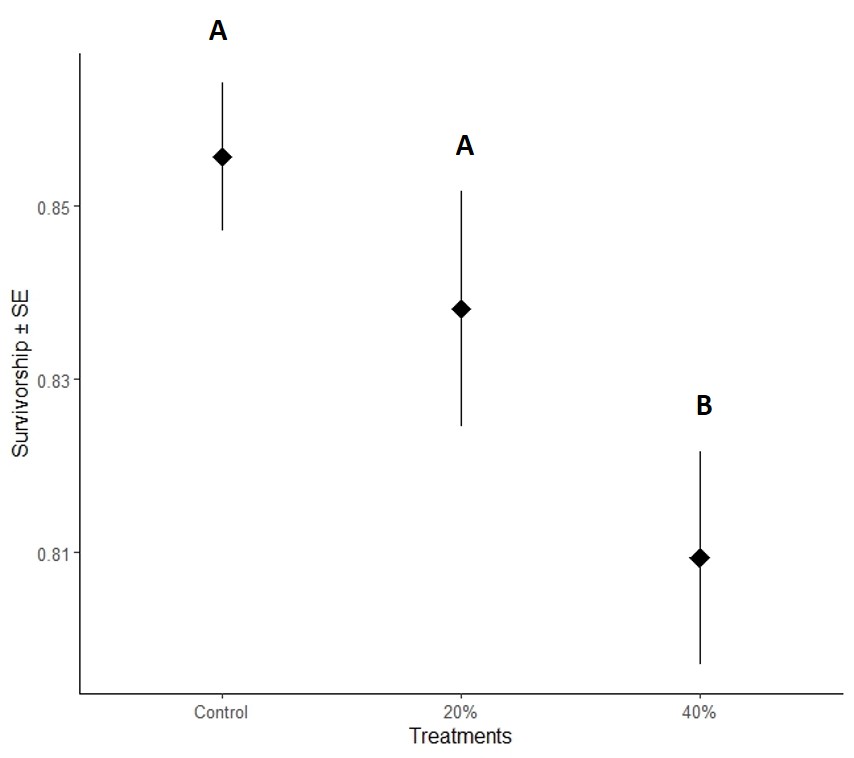

**Figure S3**: **Effect of larval diet treatment on Survivorship.** The effect of adult density on egg-to-adult survivorship. The plots represent data from all three statistical blocks. Treatments not sharing common alphabets are significantly different.

**Table S7: Model structures of the analyses in the Experiment 1, 2, 3 and 4.** The actual codes for each analysis are mentioned. “Treatment’ refer to the source patch condition.

**Experiment 1 and 4**

Model=glmer(cbind(Dispersal,Trail)~Treatment*Sex+(1|Block:Treatment:Sex)+(1|Block:Treatment)+(1|Block:Sex)+(1|Block) ,family=binomial(link="logit"),data = data)

**Experiment 2 and 3**

Model=glmer(cbind(Dispersal,Trail)~Treatment +(1|Block:Treatment)+(1|Block) ,family=binomial(link="logit"), data = data)
